# Classifying native versus foreign speech perception from EEG using linguistic speech features

**DOI:** 10.1101/2024.04.11.589055

**Authors:** Corentin Puffay, Jonas Vanthornhout, Marlies Gillis, Pieter De Clercq, Bernd Accou, Hugo Van hamme, Tom Francart

**Affiliations:** KU Leuven, Dept. Neurosciences, ExpORL, Leuven, Belgium; KU Leuven, Dept. of Electrical engineering (ESAT), PSI, Leuven, Belgium

**Keywords:** EEG decoding, deep learning, CNN, linguistics

## Abstract

When a person listens to natural speech, the relation between features of the speech signal and the corresponding evoked electroencephalogram (EEG) is indicative of neural processing of the speech signal. Using linguistic representations of speech, we investigate the differences in neural processing between speech in a native and foreign language that is not understood. We conducted experiments using three stimuli: a comprehensible language, an incomprehensible language, and randomly shuffled words from a comprehensible language, while recording the EEG signal of native Dutch-speaking participants. We modeled the neural tracking of linguistic features of the speech signals using a deep-learning model in a match-mismatch task that relates EEG signals to speech, while accounting for lexical segmentation features reflecting acoustic processing. The deep learning model effectively classifies languages. We also observed significant differences in tracking patterns between comprehensible and incomprehensible speech stimuli within the same language. It demonstrates the potential of deep learning frameworks in measuring speech understanding objectively.

## 1 Introduction

Electroencephalography (EEG) is a non-invasive method that can be used to study brain responses to sounds. Traditionally, unnatural periodic stimuli are presented to listeners, and the recorded EEG signal is averaged to obtain the resulting brain response and to enhance its stimulus-related component (Picton et al., 2005). These stimuli do not reflect everyday human natural speech, as they are repetitive, and not continuous. To investigate how the brain processes realistic speech, it is common to model the transfer function between the presented speech and the resulting brain response (Ding and Simon, 2012; Crosse et al., 2016). Such models capture the time-locking of the brain response to certain features of speech, often referred to as neural tracking. Three main model types are being used to measure the neural tracking of speech: (1) a linear regression model that reconstructs speech from EEG (backward modeling); (2) a linear regression model that predicts EEG from speech (forward modeling); and (3) classification tasks that associate synchronized segments of EEG and speech among multiple candidate segments (de Cheveigné et al., 2021; Défossez et al., 2023; Puffay et al., 2023a). For forward and backward models, the correlation between the ground truth and predicted/reconstructed signal provides the measure of neural tracking, while for the classification task, classification accuracy is utilized.

To investigate how the brain processes speech, research has focused on different features of speech signals, which are known to be processed at different stages along the auditory pathway. Three main classes have hence been investigated:

- Acoustics (e.g., spectrogram, speech envelope (Ding and Simon, 2012), f0 (Van Canneyt et al., 2021; Puffay et al., 2022))
- Lexical segmentation features (e.g., phone onsets, word onsets, (Di Liberto et al., 2015; Lesenfants et al., 2019))
- Linguistics (e.g., phoneme surprisal, word frequency, (Gillis et al., 2022; Brodbeck et al., 2018; Broderick et al., 2018; Weissbart et al., 2019; Koskinen et al., 2020; Puffay et al., 2023b))

As opposed to neural tracking studies using broad features that carry mostly acoustic information, we here select linguistic features to narrow down our focus to speech understanding. Linguistic features of speech reflect information carried by a word or a phoneme, and their resulting brain response can be interpreted as a marker of speech understanding (Gillis et al., 2022; Brodbeck et al., 2018). Considering the correlation between feature classes (Daube et al., 2019), many studies accounted for the acoustic and lexical segmentation components of linguistic features (Gillis et al., 2022; Brodbeck et al., 2018), while others did not (e.g., Broderick et al., 2018; Weissbart et al., 2019), potentially measuring the neural tracking of non-linguistic information.

Although the dynamics of the brain responses are known to be non-linear, most of the studies investigating neural tracking relied on linear models, which is a crude simplification. Later research attempted to introduce non-linearity, using deep neural networks. Such architectures relied on simple fully connected layers (de Taillez et al., 2020), recurrent layers (Monesi et al., 2020; Accou et al., 2023), or even recently transformer-based architectures (Défossez et al., 2023). For a global overview of EEG-based deep learning studies see Puffay et al. (2023a).

Most deep learning work used low-frequency acoustic features, such as the Mel spectrogram, or the speech envelope (e.g., Accou et al., 2023; Bollens et al., 2022), or higher frequency features such as the fundamental frequency of the voice, f0 (Puffay et al., 2022; Thornton et al., 2023) to improve the decoder’s performance. Although studies using invasive recording techniques showed the encoding of multiple linguistic features (Keshishian et al., 2023), very few EEG-based deep learning studies involved linguistic features (e.g., Défossez et al., 2023). In a previous study (Puffay et al., 2023b), we used a deep learning framework and measured *additional* neural tracking of linguistic features over lexical segmentation features in young healthy native Dutch speakers who listened to Dutch stimuli. This finding emphasized that a component of neural tracking corresponds to the phoneme or word rate, while another corresponds to the semantic context reflected in linguistic features. In addition, linear modeling studies (Gillis et al., 2023; Verschueren et al., 2022) suggested the relationship between understanding and the added value of linguistics. Gillis et al. (2023) used two incomprehensible language conditions (i.e. Frisian language and random-word-shuffling of Dutch speech) to manipulate speech understanding. However, within our deep learning framework, no investigations have been conducted on language data incomprehensible to the test subject.

In this article, we aim to investigate the impact of language understanding on the neural tracking of linguistics using our above-mentioned deep learning framework. Therefore, we fine-tune and evaluate our previously published deep learning framework to measure the added value of linguistics over lexical segmentation features on the neural tracking of three different stimuli: (1) Dutch, (2) Frisian, and (3) scrambled Dutch words. Additionally, we evaluate our model on a language classification task to explore whether our CNN can learn language-specific brain responses.

## 2 Methods

### 2.1 Dataset

We use the dataset from Gillis et al. (2023), which consists of 26 young native Dutch participants listening to a comprehensible story in Dutch, a list of scrambled words in Dutch, and an incomprehensible story in Frisian. The three stories were narrated by the same male native Dutch speaker, who learnt Frisian as a second language. The Dutch story is derived from a podcast series about crime cases, and the Frisian story is a translation of the Dutch. Frisian is a language related to Dutch but poorly understood by Dutch native participants who have no prior knowledge of it. The list of scrambled words consists of randomly shuffled words from the Dutch story. The duration of the three stories, from now on referred to as “language conditions”, are 20.6, 16.9, and 14.1 min. For more details, see Gillis et al. (2023).

For the pre-training of our model, we use an additional dataset from Puffay et al. (2023b), containing EEG of 60 young healthy native Dutch participants listening to 8 to 10 audiobooks of 14 min each.

### 2.2. Speech features

This study relates 4 features from EEG signals only including the linguistic features that showed a benefit over lexical segmentation features (Puffay et al., 2023b).

The investigated lexical segmentation features are: the onset of any phoneme (PO) and of any word (WO). We then tested the added value of the following linguistic features:

- Cohort entropy (CE), over PO
- Word frequency (WF), over WO

on our model’s performance, measuring the neural tracking of speech.

Example phoneme-level features are depicted in Figure 1a, and word-level features in Figure 1b.

**Figure 1.**
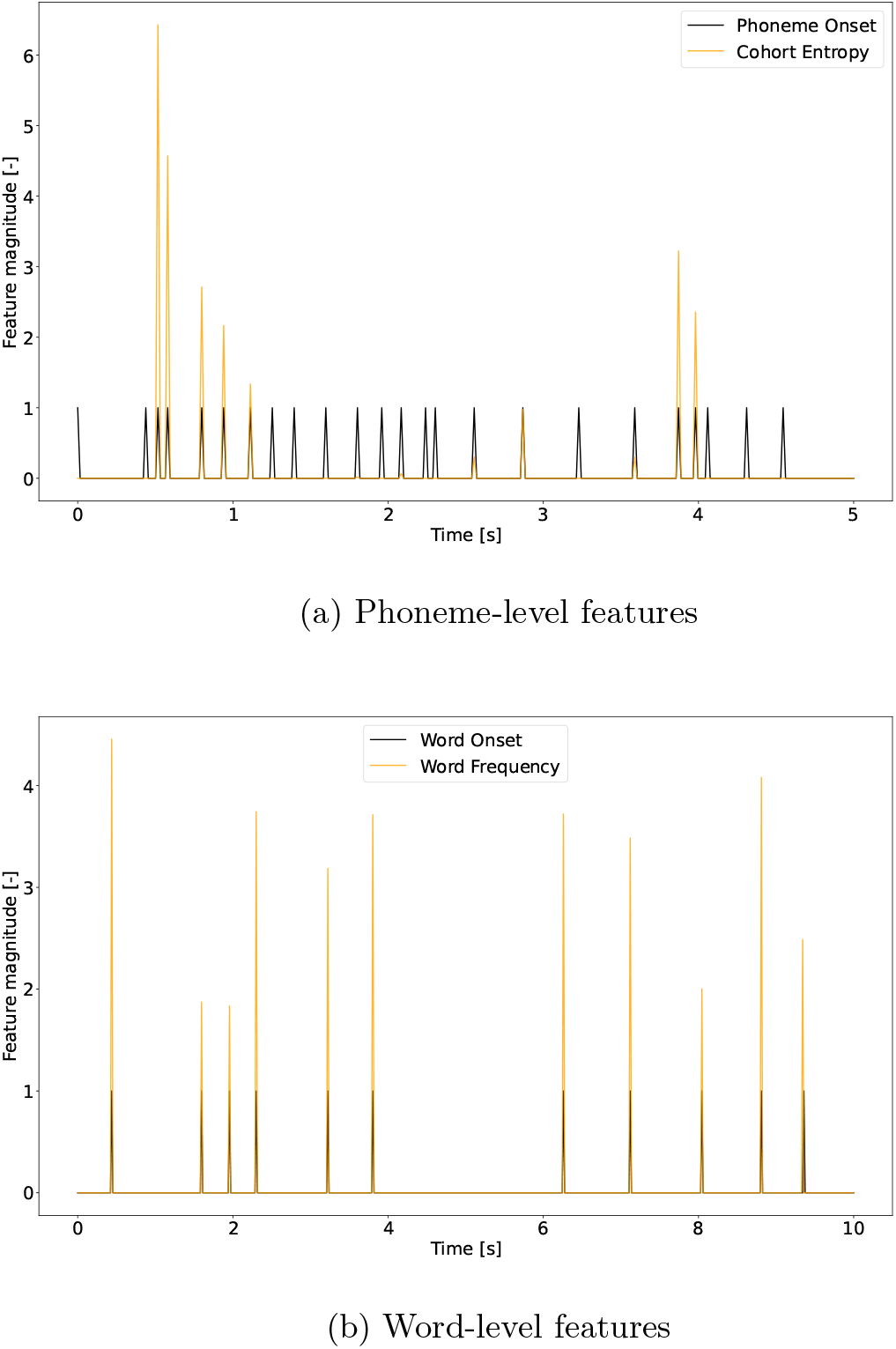
Visualization of word- and phoneme-level lexical segmentation and linguistic features. (a) Cohort entropy is depicted in yellow, phoneme onset in black, over a 5 s window. (b) Word frequency is depicted in yellow, word onset in black, over a 10 s window.

#### Lexical segmentation features

Time-aligned sequences of phonemes and words were extracted by performing a forced alignment of the identified phonemes (Duchateau et al., 2009). PO and WO are the resulting one-dimensional arrays with pulses on the onsets of, respectively, phonemes and words. Silence onsets were set to 0 for both phonemes and words.

#### Active cohort of words

Before introducing cohort entropy, the active cohort of words must be defined. Following previous studies’ definition (Brodbeck et al., 2018; Gillis et al., 2022), it is a set of words that starts with the same acoustic input at any point in the word. For example, should we find cohorts in English, the active cohort of words for the phoneme */n/* in “ban” corresponds to the ensemble of words that exist in that language starting with “ban” (e.g., “banned”,”bandwidth” etc.). For each phoneme, the active cohort was determined by taking word segments that started with the same phoneme sequence from the lexicon.

#### Lexicon

For the Dutch language, the lexicon for determining the active cohort was based on a custom word-to-phoneme dictionary (9082 words). As some linguistic features are based on the word frequency in Dutch, the prior probability for each word was computed, based on its frequency in the SUBTLEX-NL database (Keuleers et al., 2010).

For the Frisian language, the word-to-phoneme dictionary (75036 words) and the word frequencies were taken from Yılmaz et al. (2016).

#### Cohort entropy

CE reflects the degree of competition among possible words that can be created from the active cohort including the current phoneme. It is defined as the Shannon entropy of the active cohort of words at each phoneme as explained in Brodbeck et al. (2018) (see Equation 1). *CE*_*i*_ is the entropy at phoneme *i* and *p*_*word*_ is the probability of the given word in the language. The sum iterates over words from the active cohort *cohort*_*i*_.

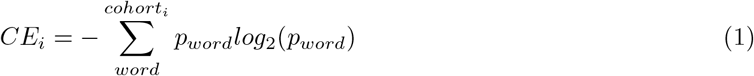

#### Word frequency

For the Dutch language, the prior probability for each word was based on its frequency in the SUBTLEX-NL database (Keuleers et al., 2010). Values corresponding to words not found in the SUBTLEX-NL were set to 0.

For the Frisian language, the word probabilities were used from Yılmaz et al. (2016). WF is a measure of how frequently a word occurs in the language and is defined in Equation 2.

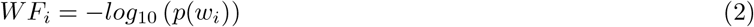

More details about their implementation can be found in previous studies (Gillis et al., 2022; Brodbeck and Simon, 2020; Puffay et al., 2023b).

### 2.3. Preprocessing

The EEG was initially downsampled using an anti-aliasing filter from 8192 Hz to 128 Hz to decrease the processing time. A multi-channel Wiener filter (Somers et al., 2018) was then used to remove eyeblink artifacts, and re-referencing was performed to the average of all electrodes. The resulting signal was band-pass filtered between 0.5 and 25 Hz using a Least Squares filter of order 5000 for the high-pass filter, and 500 for the low-pass filter, with 10% transition bands (transition of frequencies 10% above the lowpass filter and 10% below the highpass) and compensation for the group delay. We then downsampled the signal to 64 Hz.

As lexical segmentation and linguistic features are discrete, they were calculated as defined in Section 2.2, and further downsampled to 64 Hz.

### 2.4. The match-mismatch task

In this study, we use the performance of the match-mismatch (MM) classification task (de Cheveigné et al., 2021) to measure the neural tracking of different speech features (Figure 2). We use the same paradigm as Puffay et al. (2023b). The model is trained to associate the EEG segment with the matched speech segment among two presented speech segments. The matched speech segment is synchronized with the EEG, while the mismatched speech segment occurs 1 second after the end of the matched segment. These segments are of fixed length, namely 10 s for word-based features and 5 s for phoneme-based features, to provide enough context to the models as hypothesized by Puffay et al. (2023b). This task is supervised since the matched and mismatched segments are labeled. The evaluation metric is classification accuracy.

**Figure 2.**
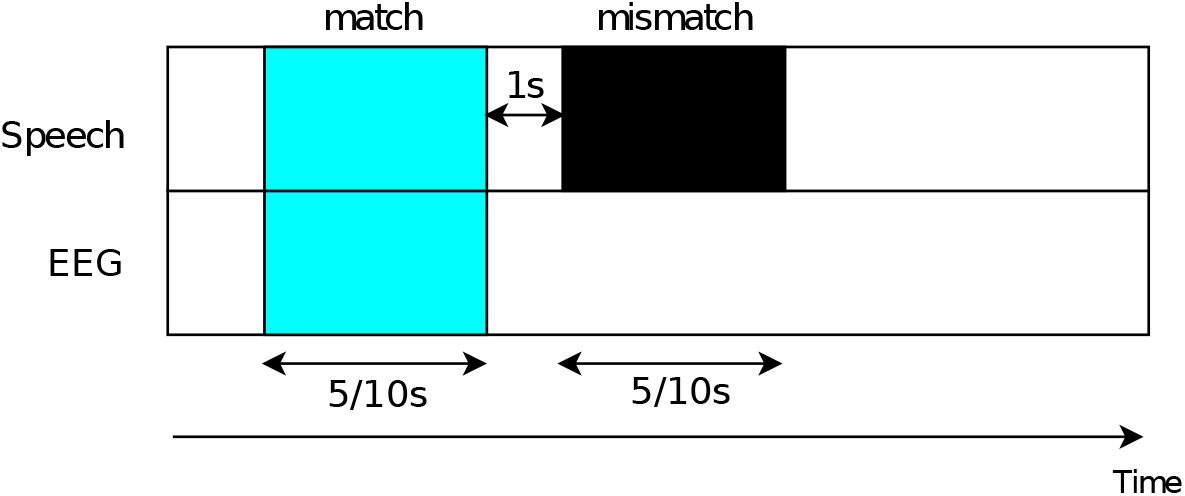
Match-mismatch classification task. The match-mismatch task is a binary classification paradigm that associates the herewith blue EEG and speech segments. The matched speech segment is synchronized with the EEG (blue segment), while the mismatched speech occurs 1 second after the end of the matched segment (black segment). The figure depicts segments of 5 s and 10 s, which will be lengths used in our studies for the phoneme and word levels respectively.

### 2.5. Multi-input features convolutional neural network (Puffay et al., 2023b)

In Puffay et al. (2023b), we developed a multi-input convolutional neural network (MICNN) model that aims to relate different features of the presented speech to the resulting EEG. The MICNN model has 127k parameters and is trained using binary cross-entropy as its loss function (Adam optimizer, 50 epochs, learning rate: 10^−^3).

We used early stopping as regularization. It is trained to perform well on the MM task presented in Section 2.4. Through the MM task, the MICNN model learns to measure the neural tracking of speech features, which we can thereafter use to quantify the added value of one speech feature over another. By inputting multiple features, we make sure to account for redundancies and correlations between them and enable interpretation of what information makes the model better on the MM task. In our case, our models enable us to quantify the added value of a given linguistic feature (WF or CE) over their corresponding lexical segmentation feature (WO or PO, respectively)

### 2.6. Fine-tuning and evaluation on validation datasets

We performed two fine-tuning conditions: one subject-independent (language finetuning) and one subject-dependent (subject fine-tuning). For both, we keep the training parameters mentioned in Section 2.5, and solely change the data used for training and evaluation.

For the language condition fine-tuning, we trained a separate model for each subject, including data from the other 25 subjects, for each of the three language conditions (i.e., Dutch, Frisian, and scrambled Dutch): We exclude a selected subject and separate the data from the 25 other subjects in an 80%/10%/10% training/validation/test split. For the 25 other subjects, the first and last 40% of their recording segment were used for training. The first 10% of the remaining 75% was used for validation (i.e., for regularization) and the remaining 10% to get an estimate of the accuracy of unseen speech data. Once the model is fine-tuned, we evaluate it on the selected subject.

For the subject-specific fine-tuning, a 25%/25%/50% training/validation/test split was performed. We used the validation set for regularization. The first and last 12.5% of the recording segment were used for training. The first 25% of the remaining 75% was used for validation and the remaining 50% for testing. For each set (training, validation, and testing), each EEG channel was then normalized by subtracting the mean and dividing by the standard deviation.

### 2.7. Language condition classification

Inspired by the support vector machine (SVM) utilization for aphasia classification from Clercq et al. (2024), we use the MM accuracy obtained with four models: the control (word onset or phoneme onset) and linguistic (cohort entropy or word frequency) models for both fine-tuning conditions. We chose to use only the fine-tuned conditions as the non-fine-tuned one was biased towards giving better performance on the Dutch condition. These four MM accuracy values constitute the features provided to the SVM to solve a one-vs-one classification: did the person listen to one or another selected language condition. We consider three language conditions: Dutch, scrambled Dutch, and Frisian, which in total leads to three binary classification tasks.

We used a radial basis function kernel SVM and performed a nested cross-validation approach. In the inner cross-validation, the C-hyperparameter (determining the margin) and pruning were optimized (accuracy-based) and tested in a validation set using 5-fold cross-validation. Predictions were made on the test set in the outer loop using leave-one-subject-out cross-validation. We computed the receiver operating characteristic (ROC) curve and calculated the area under the curve (AUC), and further reported the accuracy, F1-score, sensitivity, and specificity of the classifier.

## 3 Results

### 3.1. Impact of the language on the neural tracking of linguistic features

We only depict results with language or subject fine-tuning as pure evaluation would potentially give a performance advantage to the model on Dutch because of the pretraining on Dutch stimuli. We still show the non-fine-tuned results in Appendix A.

#### 3.1.1 Evaluation of the trained MICNN across languages with language fine-tuning

Overall, at both the word and the phoneme levels, the increase in neural tracking when adding the linguistics on top of lexical segmentation features is significant. We depict in Figure B1a and B1b the models performances at the phoneme and the word levels across stimuli.

At the phoneme level, the match-mismatch accuracy obtained for Dutch, Frisian, and Sc. Dutch was higher for the linguistics than for the control models (Wilcoxon signed-rank test: *p <* 0.001, *W* = 17.5, *p <* 0.01, *W* = 60, and *p <* 0.001, *W* = 25 respectively) At the word level, the match-mismatch accuracy obtained for Dutch and Sc. Dutch was higher for the linguistics than for the control models (Wilcoxon signed-rank test: *p <* 0.001, *W* = 36, and *p* = 0.016, *W* = 82 respectively), while not significantly higher for the Frisian language (Wilcoxon signed-rank test: *p* = 0.19, *W* = 36).

Figure 3 depicts for all three stimuli, the difference in the MM accuracy between L and C conditions for phoneme-level features. We observed no significant difference in the Frisian-Dutch comparison (Wilcoxon signed-rank test, *W* = 160, *p* = 0.71), Dutch-Sc. Dutch comparisons (Wilcoxon signed-rank test, *W* = 126, *p* = 0.22), and the Frisian-Sc. Dutch comparison (Wilcoxon signed-rank test, *W* = 112, *p* = 0.11).

**Figure 3.**
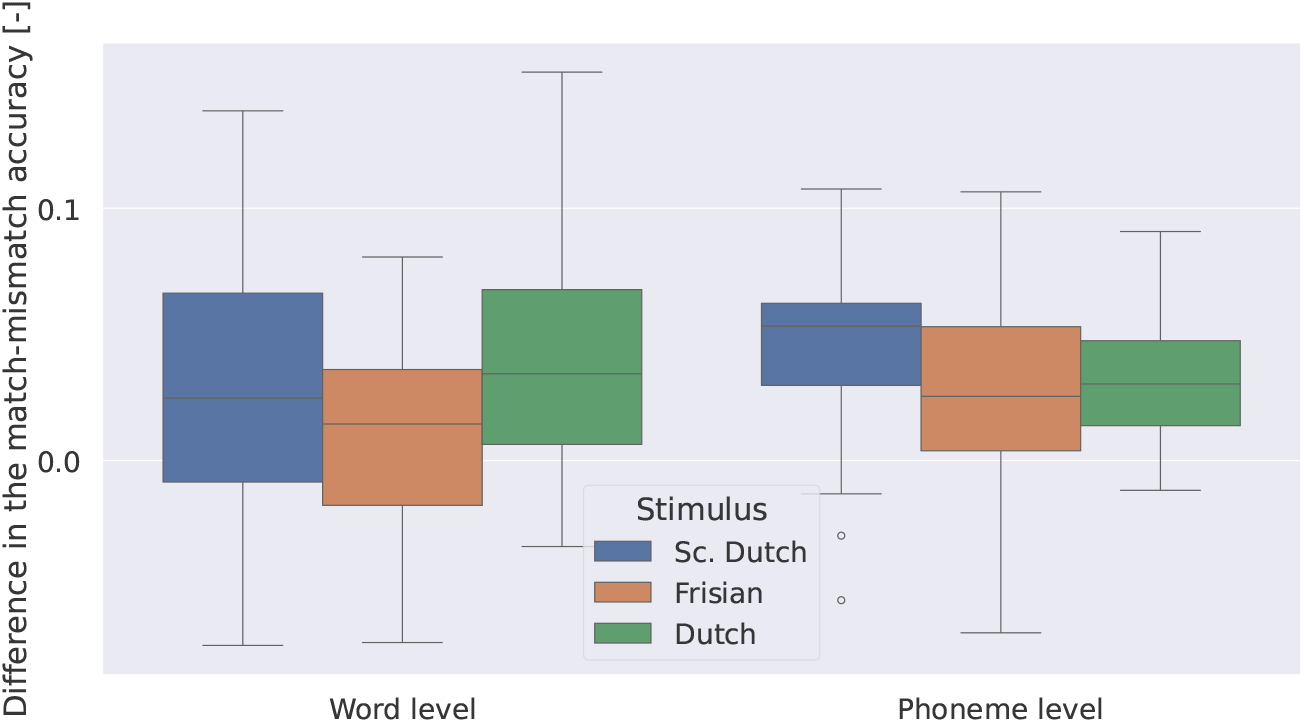
L - C accuracy for three stimuli with a language-finetuned model. L corresponds to the MM accuracy obtained by (1) the word frequency model at the word level; (2) the cohort entropy model at the phoneme level. C corresponds to the MM accuracy obtained by (1) the word onset model at the word level; (2) the phoneme onset model at the phoneme level.

**Figure 4.**
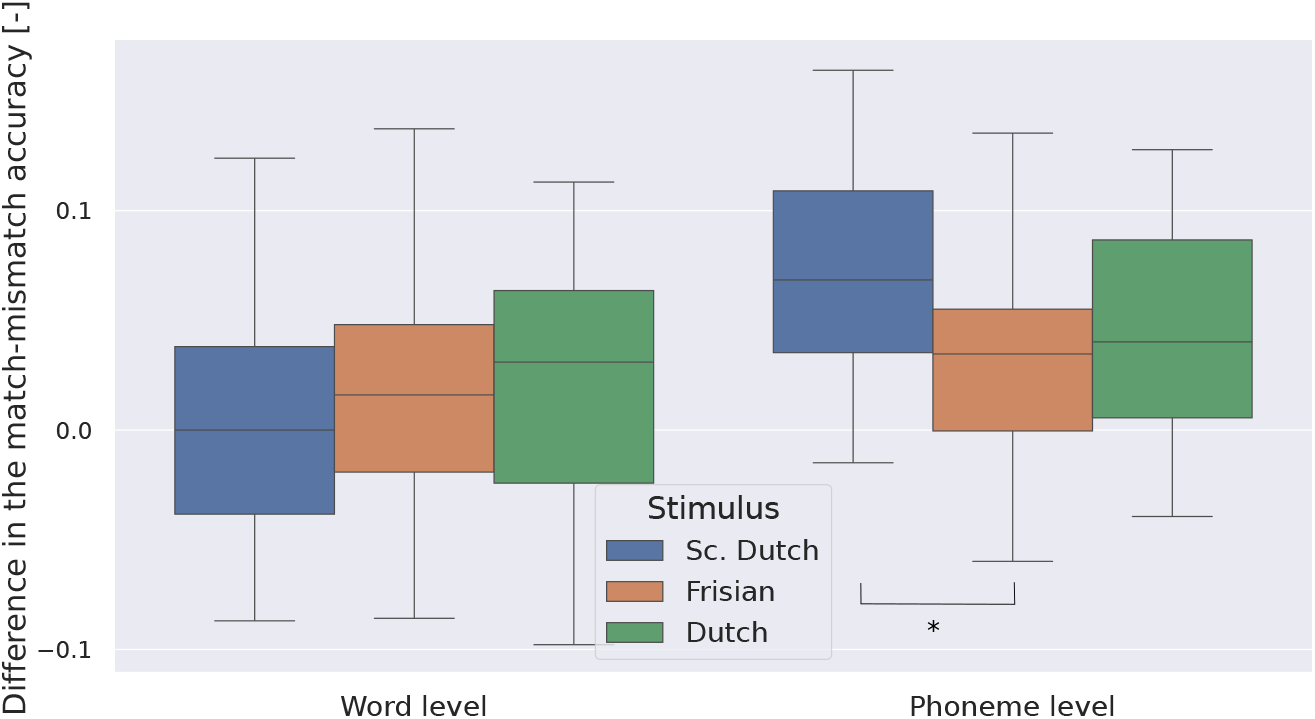
L - C accuracy across recording lengths for three stimuli with a subject-finetuned model. L corresponds to the MM accuracy obtained by (1) the word frequency model at the word level; (2) the cohort entropy model at the phoneme level. C corresponds to the MM accuracy obtained by (1) the word onset model at the word level; (2) the phoneme onset model at the phoneme level. (∗ : *p <* 0.05)

We also depict for all three stimuli, the L-C accuracy at the word level. We observed a significant increase of the L-C accuracy of Dutch over the Frisian’s (Wilcoxon signed-rank test, *W* = 86.0, *p* = 0.022), however no significant difference in the Dutch-Sc. Dutch and Frisian-Sc. Dutch comparisons (Wilcoxon signed-rank test, Dutch-Sc. Dutch: *W* = 155, *p* = 0.62, Frisian-Sc. Dutch: *W* = 145, *p* = 0.45).

To see whether the model could be improved by introducing subject information, we add a subject fine-tuning step in the next Section (for details about the method, see Section 2.6).

#### 3.1.2 Evaluation of the trained MICNN across languages with language and subject fine-tuning

We depict results up to half of the recording length of the shortest stimulus for each subject (i.e., 7.5 min) as we used the other half to fine-tune the model.

Figure 3 depicts for all three stimuli, the L-C accuracy at the phoneme level. We observed a significant increase of the L-C accuracy of Frisian over the Sc. Dutch accuracy (Wilcoxon signed-rank test, *W* = 88.0, *p* = 0.025), however no significant difference in the Dutch-Sc.-Dutch and Frisian-Dutch comparisons (Wilcoxon signed-rank test,*W* = 121, *p* = 0.17, and *W* = 149, *p* = 0.51 respectively).

We also depict for all three stimuli, the L-C accuracy at the word level. At the longest length of the shortest stimuli (i.e., 7.5 min), we observed no significant difference’ in the L-C accuracy across Dutch, Frisian, and scrambled Dutch (Wilcoxon signed-rank test, Dutch-Frisian: *W* = 159, *p* = 0.68, Dutch-Sc.Dutch: *W* = 143, *p* = 0.42, and Sc.Dutch-Frisian: *W* = 154, *p* = 0.60).

### 3.2. Language classification task

Figure 5 depicts the SVM classification results of the three binary classification tasks: (1) Dutch vs. Frisian, (2) Dutch vs. Scrambled Dutch, and (3) Frisian vs. Scrambled Dutch. For more details about the methods, see Section 2.7.

**Figure 5.**
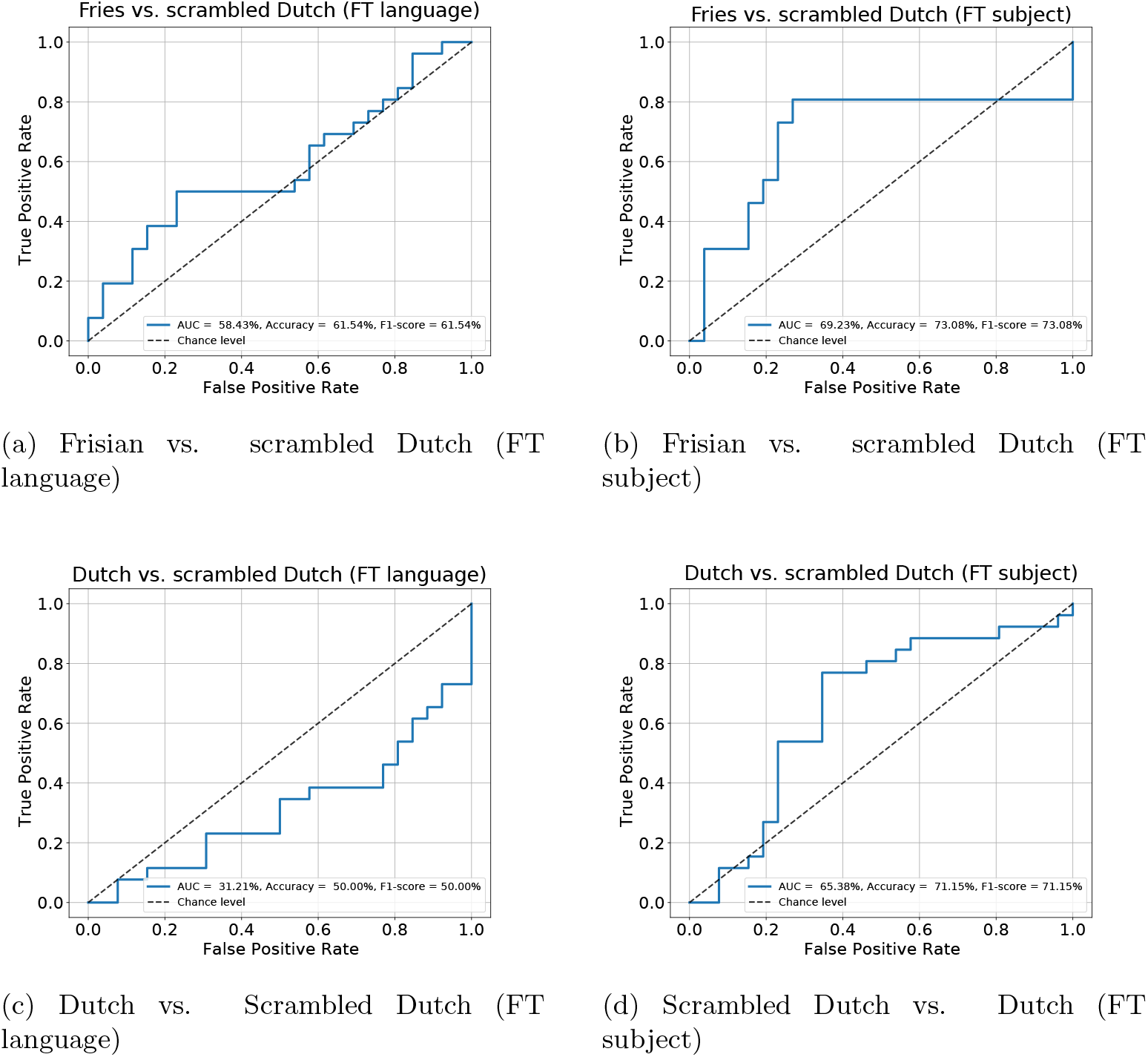
SVM performance across fine-tuning conditions. The performance is depicted for each condition as a ROC curve plotting the true positive rate as a function of the false positive rate. (a) Scrambled Dutch vs. Frisian classification with language fine-tuning; (b) with subject fine-tuning; (c) Scrambled Dutch vs. Dutch classification with language fine-tuning; (d) with subject fine-tuning.

When evaluated over all subjects, our SVM classifier correctly classified the scrambled Dutch from the Frisian condition with an accuracy of 61.5% and 73% for the FTL and FTS conditions, respectively. In addition, the classifier correctly classified the scrambled Dutch from the Dutch condition with an accuracy of 50% and 71.15% for the FTL and FTS conditions, respectively. We do not show the results for the Dutch vs. Frisian task, as the classifier performed close to the chance level.

## 4 Discussion

We evaluated a deep learning framework that measures the neural tracking of linguistics on top of lexical segmentation features in different language understanding conditions. Although we used the same dataset, direct comparison with Gillis et al. (2023) is difficult, considering the difference in the models, and the features provided to the model.

As our model was trained uniquely on Dutch before, the model might not have learned the typical brain response to Frisian linguistics, or scrambled Dutch, thus leading to overfitting on Dutch, impairing the objective measure of linguistics tracking on other language conditions. To avoid this bias, we fine-tuned our model on Frisian and scrambled Dutch data before respective evaluations. Since we are interested in the linguistics added value over lexical segmentation features, we compared the difference between the linguistic and lexical segmentation models’ performance across language conditions. For cohort entropy, although there is no significant difference between language conditions in the linguistics added value, the one of Frisian is systematically lower. For word frequency, we observed a significant increase in the added value of linguistics for Dutch over Frisian. In addition, although not significantly different, the linguistics added value also appeared lower for Frisian than scrambled Dutch. This finding suggests that a language that is not understood might show a lower linguistics added value. Regarding the scrambled Dutch results performing non-significantly different than Dutch, although the subjective rating of understanding was very low, individual words are still in Dutch and thus understood. Cohort entropy and word frequency are features that are independent of the order of words in the sentence, which might explain why we do not observe a drop in the neural tracking of linguistics.

Language processing in the brain is influenced by our memory, and top-down processing (Gwilliams and Davis, 2022; Laura Gwilliams and King, 2023), and might thus have a strong subject-specific component in the response to linguistic features. We therefore decided to fine-tune the models on each subject before evaluation on top of the language fine-tuning. The only significant difference we observed was for cohort entropy between scrambled Dutch and Frisian. We note that the subject fine-tuning diminished the data used per subject by 50% (i.e., up to 7.5 min of recording) for evaluation, which might not be sufficient to get a good estimate of the accuracy. We therefore do not interpret further the subject fine-tuning condition.

With SVM classifiers, we were able, from the match-mismatch accuracies on our different features, to classify the Frisian vs. the scrambled Dutch condition, as well as the Dutch vs. the scrambled Dutch condition. This suggests that neural tracking of linguistics and lexical segmentation features differs between continuous and scrambled speech. We expected the classifier to be able to differentiate Frisian from Dutch, as participants were not Frisian speakers. Our hypotheses to explain this phenomenon are fourfold: (1) Frisian is a language that is too similar to Dutch to measure a difference in linguistic tracking, therefore there is some understanding by the participants, as emphasized by the subjective rating from Gillis et al. (2023) ^*‡*^; (2) Linguistic and lexical segmentation features are too correlated, notably because their only differ in the magnitude, which might be too limited to describe the language complexity (3) the magnitude of linguistics has a distribution that tends to be skewed towards the value 1 (i.e., the magnitude of lexical segmentation features) in our three stimuli (see Gillis et al. (2023)). A more controlled speech content (e.g., sentences with uncommon words) might make the impact of linguistics larger; (4) An additional concern can be added for word frequency: the most frequent words in the language are non-content words (e.g., “and”, “or”, “of”). In addition, most of these words are short words. The model might therefore have learnt a spurious content vs. non-content words threshold from the word frequency, which can be globally narrowed-down to short vs. long words. The length of words can possibly be derived from the word onsets as well. The model could therefore simply use word onset information and omit the magnitude provided by linguistics, which would explain the low benefit of adding word frequency over word onsets.

A possible shortcoming of our training paradigm is the use of a single language for pre-training (i.e., Dutch), which might provide the fine-tuning insufficient abilities to generalize to other language conditions. To solve this issue and preserve a necessary pre-training step for complex deep learning frameworks, we could change our experimental paradigm by: (1) keeping the same language across understanding conditions to avoid biasing the model during pre-training; and (2) avoid random word-shuffling to preserve the word context in sentences. Other non-understanding conditions could involve vocoded speech or degraded-SNR speech as done in Accou et al. (2021). We also evaluated our framework to a speech rate paradigm (Verschueren et al., 2022). However, although we observed a decreased neural tracking of linguistics in challenging listening scenarios (i.e., very high speech rates), we also observed an equivalent decreased neural tracking of lexical segmentation features. We could thus not draw any conclusions whether the nature of this decrease was acoustic or linguistic. Another pitfall in our comparison across languages is that our linguistic features both rely on word frequencies. The word frequency values were therefore calculated for Dutch and Frisian, respectively. Our participants being Dutch speakers not speaking Frisian, have an language representation in the brain corresponding to the Dutch word frequencies, and not to the Frisian one. This might thus result in a lower neural tracking of linguistics when listening to Frisian content compared to Dutch content. Linguistic features, as we use them now, are very constrained: they mainly give information about the word or phoneme frequency in the language. Language models are known to capture more information. As an example, the Bidirectional Transformers for Language Understanding (i.e., BERT) (Devlin et al., 2019) model carries phrase-level information in its early layers, surface (e.g., sentence length) and syntactic (e.g., word order) information in the intermediate layers and semantic features (e.g., subject-verb agreement) in the late layers (Jawahar et al., 2019). Such representations could contain more detailed information about the language than our current linguistic features. Défossez et al. (2023) used larger pre-trained speech encoder models, and following up on this work, we could use language model layers, providing information about the structure of language that can be related to brain responses (Goldstein et al., 2022; Caucheteux and King, 2022).

## 5 Conclusion

In this article, we investigated the impact of language understanding on the neural tracking of linguistics. We demonstrated that our previously developed deep learning framework can classify language using the neural tracking of linguistics. We explored the ability and the limitations of state-of-the-art linguistic features to objectively measure speech understanding using lexical segmentation features as our acoustic tracking baseline. Our findings along with the current literature support the idea that, considering this framework, further work should be dedicated to (1) designing new linguistic features using recent powerful language models, and (2) using incomprehensible and comprehensible speech stimuli from the same language, to facilitate the comparison across conditions.

## 6. Acknowledgements

The authors thank all the participants for the recordings, as well as Wendy Verheijen, Marte De Jonghe, Kyara Cloes, Amelie Algoet, Jolien Smeulders, Lore Kerkhofs, Sara Peeters, Merel Dillen, Ilham Gamgami, Amber Verhoeven, Lies Bollens, Vitor Vasconcelos and Amber Aerts for their help with data collection. Funding was provided by FWO fellowships to Bernd Accou (1S89622N), Marlies Gillis (1SA0620N; additional Internal Funds KU Leuven: PDMT1/23/011), Corentin Puffay (1S49823N), and Jonas Vanthornhout (1290821N).

## Appendix A. Evaluation of the trained MICNN across languages without fine-tuning

In this section, we use the MICNN model from our previous study (Puffay et al., 2023b), without fine-tuning.

Figure A1a depicts the MM accuracy obtained for each subject using control (C) and linguistic (L) models across language conditions at the phoneme level. At the group level, no significant differences were found between C and L conditions for both Frisian and scrambled Dutch. As expected from previous findings, we found a significant difference between C and L conditions for Dutch (Wilcoxon signed-rank test, *W* = 41, *p <* 0.001). We also depict the difference between L and C models’ accuracy across language conditions in Figure A1b.

Figure A2a depicts the MM accuracy obtained for each subject using control (C) and linguistic (L) models across language conditions at the word level. At the group level, no significant differences were found between C and L conditions for both Dutch and scrambled Dutch (Wilcoxon signed-rank test, *W* = 104, *p* = 0.071 and *W* = 119, *p* = 0.165 respectively). Unexpectedly, we found a significant decrease of L compared to C conditions for Frisian (Wilcoxon signed-rank test, *W* = 34, *p* = 1.09 ∗ 10^−4^). We also depict the difference between L and C models’ accuracy across language conditions in Figure A2b.

Please note that the model was trained on Dutch stimuli, which might have introduced a bias to better model the linguistics benefit over the lexical segmentation features in Dutch and not in Frisian. In addition, evaluating models on different amounts of data (the three stimuli are of different duration) might be unfair for comparison. In the next step, we, therefore, fine-tune the model (for details about the method, see Section 2.6) for each language and evaluate it on the maximum length of the shortest stimuli (i.e., 14.1 min).

## Appendix B. Neural tracking of linguistic over control models across languages

**Figure A1:**
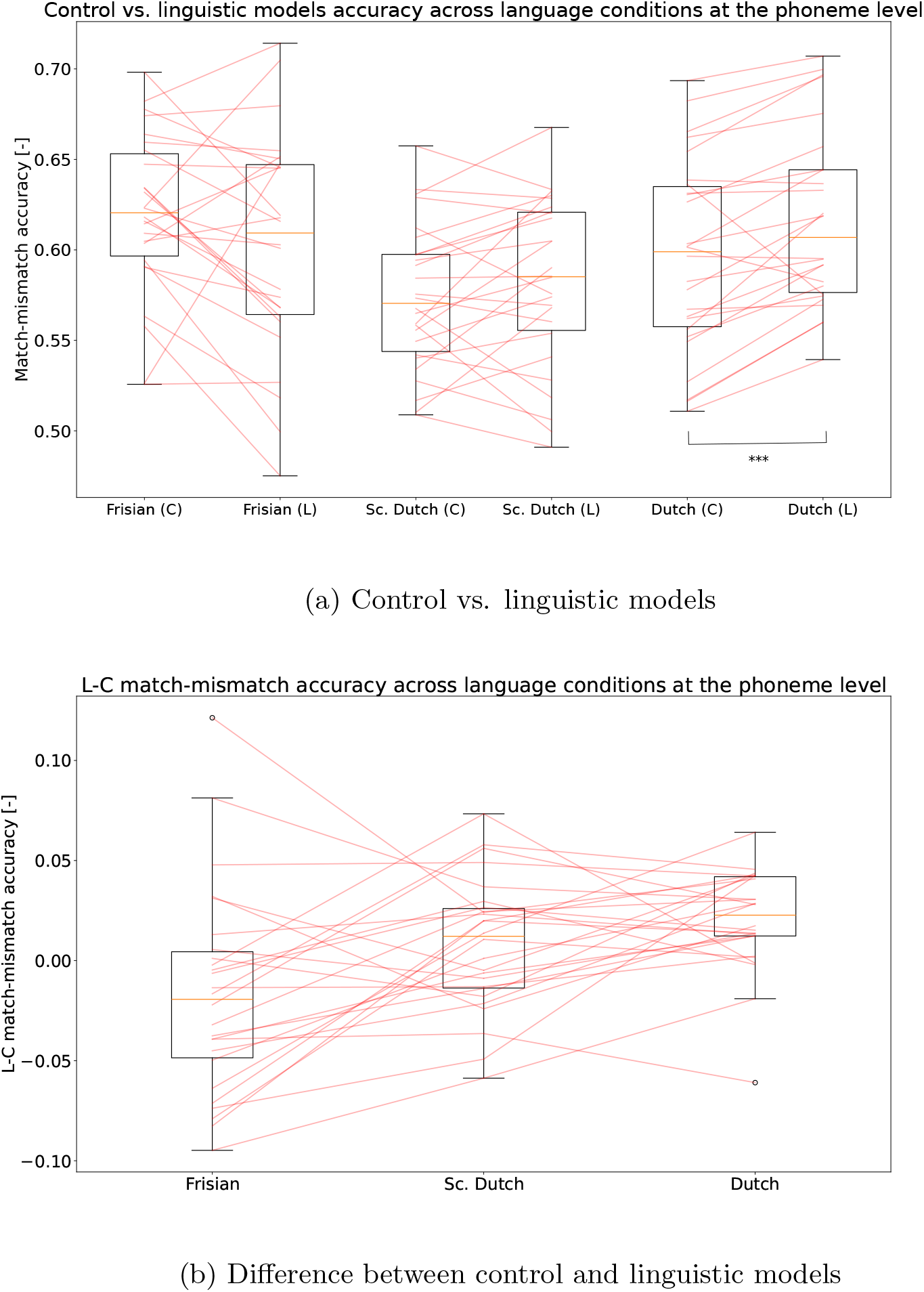
Control (C) vs. linguistic (L) models’ accuracy across language conditions at the phoneme level. The boxplots represent the accuracy obtained per subject, and the red lines connect subjects between (a) the performance of the C and L model for a given participant; (b) the performance’s difference on a language condition and another. The language conditions are Frisian, scrambled Dutch (Sc. Dutch), and Dutch. *Wilcoxon signed-rank test:* (∗ ∗ ∗ : *p <* 0.001)

**Figure A2:**
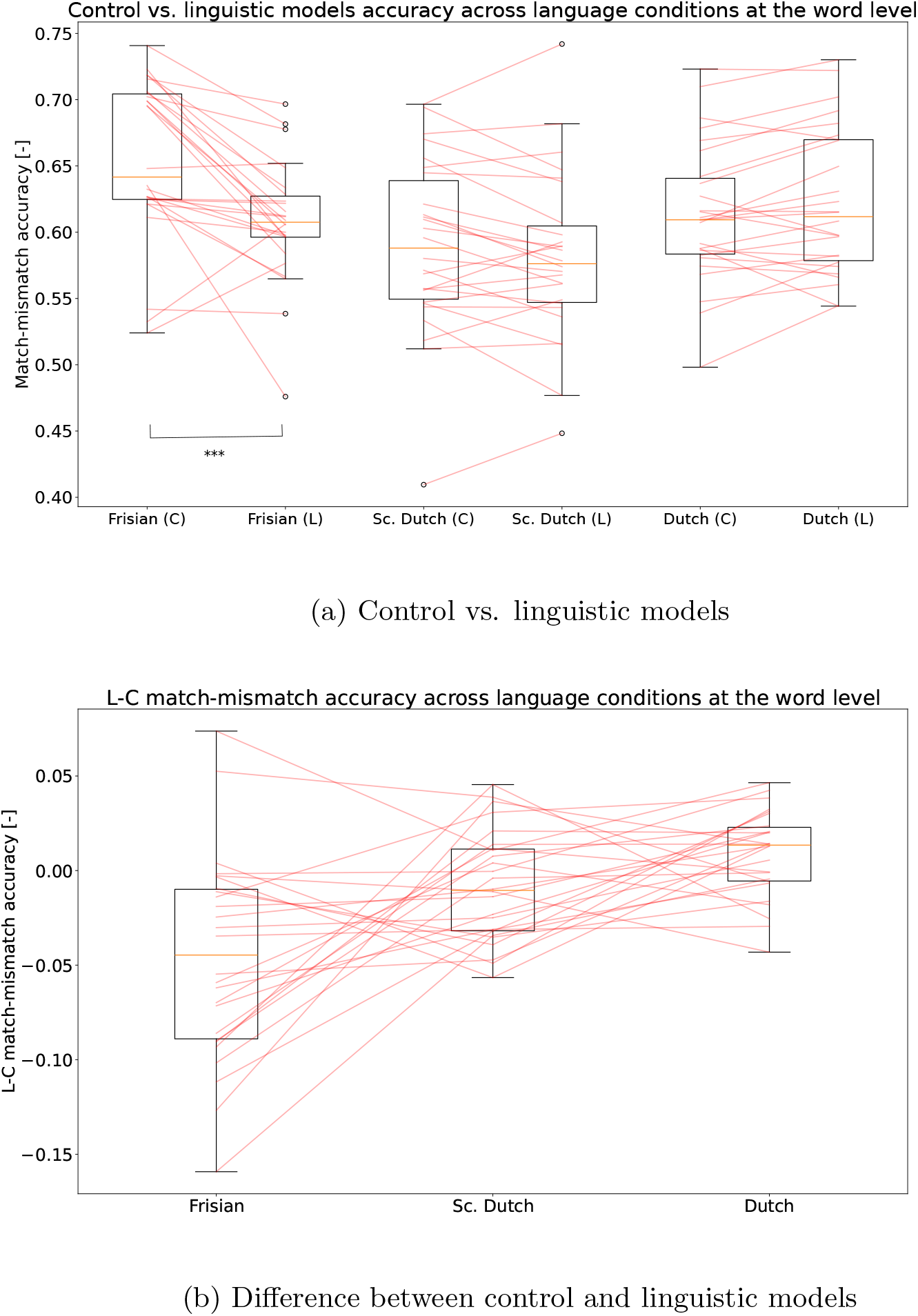
Control (C) vs. linguistic (L) models’ accuracy across language conditions at the word level. The boxplots represent the accuracy obtained per subject, and the red lines the slope between language conditions for a given subject. (a) The performance of the C and L models across language conditions; (b) L-C performance. The language conditions are Frisian, scrambled Dutch (Sc. Dutch), and Dutch. *Wilcoxon signed-rank test:*(∗ ∗ ∗ : *p <* 0.001)

**Figure B1:**
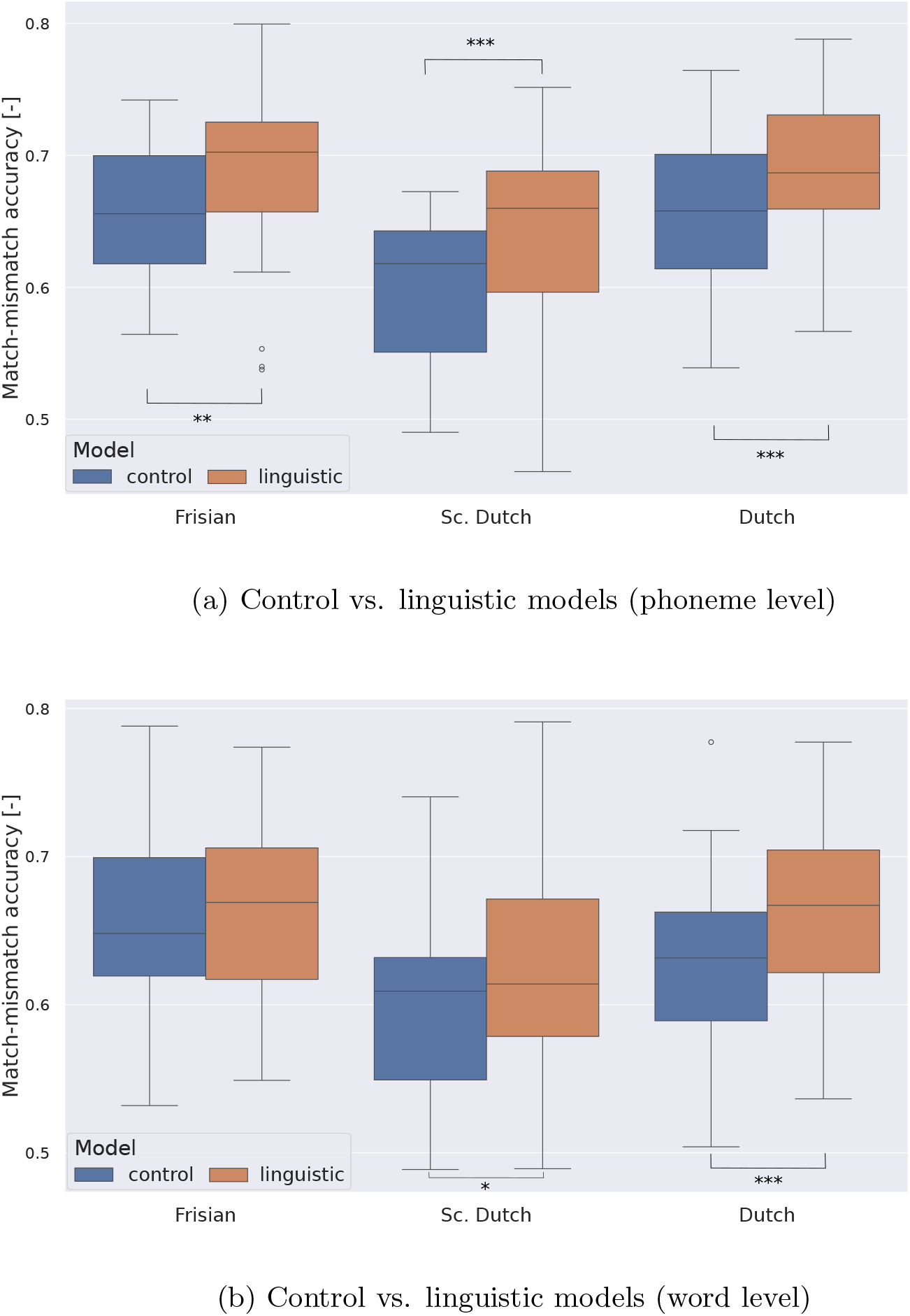
Control (C) vs. linguistic (L) models’ accuracy across language conditions at the phoneme and word level in the language fine-tuning condition. The boxplots represent the accuracy obtained per subject at (a) the phoneme level (b) the word level. The language conditions are Frisian, scrambled Dutch (Sc. Dutch), and Dutch. *Wilcoxon signed-rank test:* (∗ : *p <* 0.05, ∗∗ : *p <* 0.01, ∗ ∗ ∗ : *p <* 0.001)

## Footnotes

‡ From Gillis et al. (2023), we know that the speech understanding median subjective rating for the Dutch condition was 100%, while the Frisian and scrambled Dutch were 50% and 10.5%, respectively. The value for Frisian is strangely high and we believe it might in reality be lower.

